# The prophase oocyte nucleus is a homeostatic G-actin buffer

**DOI:** 10.1101/2020.10.30.353961

**Authors:** Kathleen Scheffler, Federica Giannini, Binyam Mogessie

**Author notes:** Equal contribution authors (Kathleen Scheffler, Federica Giannini).

## Abstract

Formation of healthy mammalian eggs from oocytes requires specialised F-actin structures. F-actin disruption produces aneuploid eggs, which are a leading cause of human embryo deaths, genetic disorders, and infertility. We found that oocytes regulate F-actin organisation and function by promptly transferring excess monomeric G-actin from the cytoplasm to the nucleus. Inside healthy oocyte nuclei, transferred monomers form dynamic F-actin structures, a conserved feature that significantly declines with maternal age. Monomer transfer must be controlled tightly. Blocked nuclear import of G-actin triggers assembly of a dense cytoplasmic F-actin network, while excess G-actin in the nucleus dramatically stabilises nuclear F-actin. Imbalances in either direction predispose oocytes to aneuploidy. The large oocyte nucleus is thus a homeostatic G-actin buffer that is used to maintain cytoplasmic F-actin form and function.

**One Sentence Summary:** Mammalian oocyte nuclei buffer cytosolic G-actin

---

Mammalian eggs are formed when oocyte chromosomes are segregated during meiosis [1], successful completion of which is a prerequisite for healthy embryogenesis and development. Meiotic errors in oocytes are a leading cause of aneuploidies that underlie human infertility and genetic disorders such as Down’s syndrome [2]. Distinct F-actin polymers assembled from soluble G-actin monomers ensure the production of healthy eggs from oocytes. These actin-rich drivers include a network of cytoplasmic actin filaments which oversee long-range vesicle transport [3] and asymmetric division in mammalian oocytes [4–6]. In addition, oocyte meiotic spindles in several mammalian species contain actin filaments that aid microtubule fibres in chromosome separation [7].

We have now found that the intact nucleus of prophase-arrested, non-manipulated mouse oocytes also contains prominent actin filaments (Fig. 1A). Using fluorescent phalloidin, we detected actin filaments and bundles in the nucleoplasm of 80% of fixed oocytes that we analysed (Fig. 1, B and C). Notably, this observation was strain- and species-independent as we could detect nuclear F-actin structures in oocytes isolated from outbred and inbred mouse (Fig. S1, A and B) and sheep ovaries (Fig. S1C). Super-resolution live imaging of nuclear F-actin using very low and non-stabilising concentrations of a fluorescently-labelled actin nanobody (nuclear actin chromobody)[8] (Fig. S1D) further revealed that these filaments are highly mobile – filaments continuously move about in non-directed fashion within the nucleoplasm (Fig. 1D and movies S1 and S2). Nuclear F-actin presence remarkably associated with distinct organisation of chromatin surrounding the nucleolus (Fig. S1, E and F), a marker of high oocyte meiotic competence and developmental capacity [9]. This indicates that nuclear F-actin structures are a common feature of healthy mammalian oocytes.

**Fig. 1.**
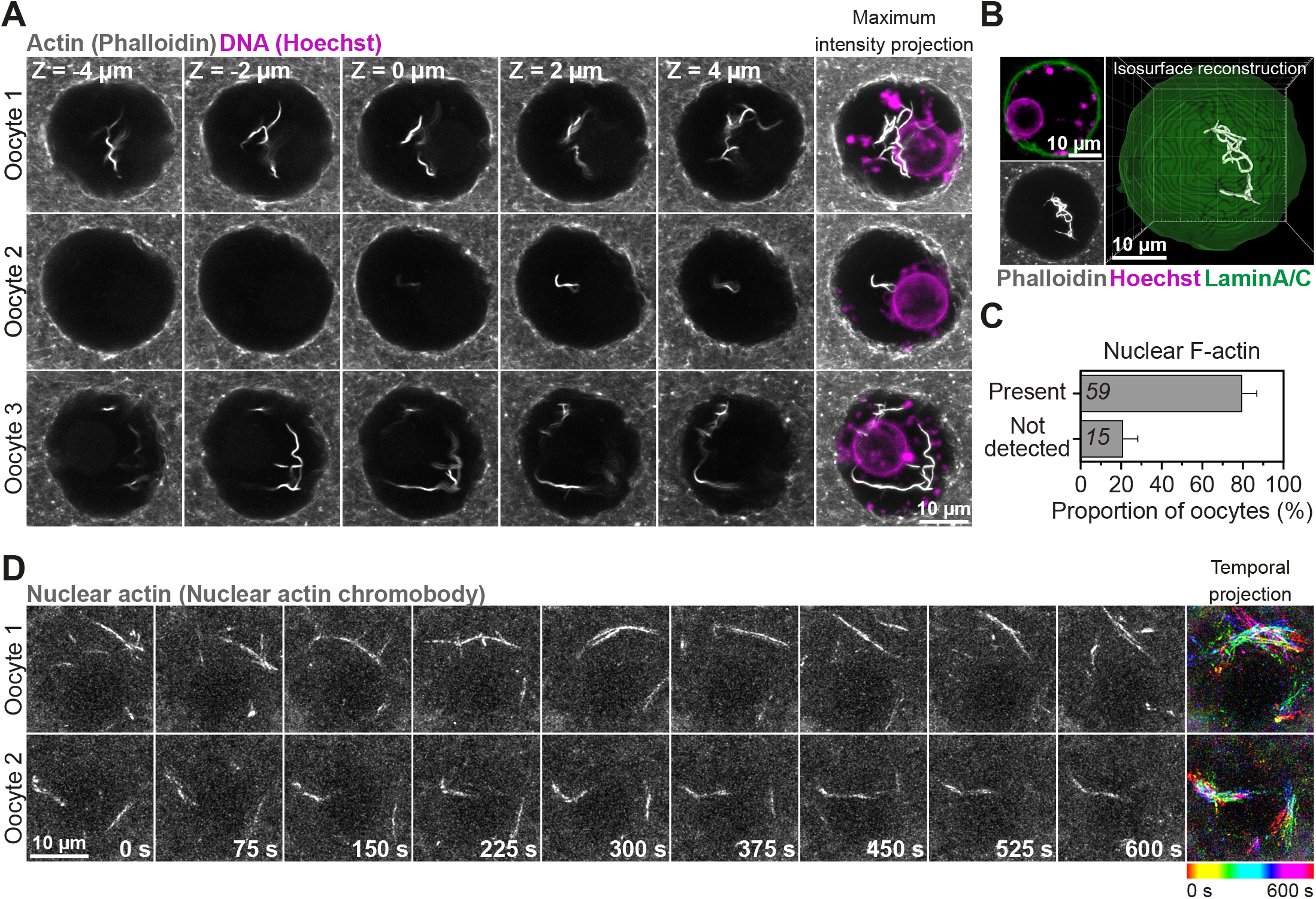
Mammalian prophase oocyte nuclei contain prominent actin filaments. (**A**) Phalloidin labelled nuclear actin filaments (grey) and chromosomes (Hoechst, magenta) in three prophase-arrested mouse oocytes. Single confocal sections spaced 2 μm apart and corresponding maximum intensity projections are shown. (**B**) Pipeline for three-dimensional isosurface reconstruction nuclear membrane (green) and nuclear F-actin (grey). DNA is shown in magenta. (**C**) Quantification of nuclear F-actin presence in prophase-arrested mouse oocytes. Data are from 3 independent experiments. (**D**) Super-resolution live imaging of nuclear actin filaments in two prophase-arrested mouse oocytes. Nuclear F-actin is labelled using non-stabilising concentration of a fluorescent nanobody (nuclear actin chromobody). Color-coded temporal projection images indicate continuous mobility of filaments in the 600 seconds observation time.

Unexpectedly, we observed that disruption of the oocyte cytoplasmic actin network using Cytochalasin D [3] triggers excessive nuclear F-actin assembly (Fig. 2, A and B). To confirm this, we imaged by high-resolution microscopy fluorescent phalloidin-labelled nuclear actin filaments, then selectively reconstructed them in three-dimensions and quantified their volume inside the nuclei (marked with nuclear membrane antibodies) of DMSO (Control)-or Cytochalasin D-treated oocytes (Fig. 2A). This showed a near forty-fold increase in nuclear F-actin volume after disruption of the cytoplasmic actin network (Fig. 2C). Bulk transfer of G-actin from a large cytoplasm to a smaller nuclear volume in Cytochalasin D-treated oocytes could increase nuclear actin monomer concentration and cause excess F-actin polymerisation. Such filaments are likely to be drug-resistant because Cytochalasin D is generally less effective at high actin monomer concentration [10]. We tested this possibility using Latrunculin, a mechanistically distinct compound that complements our Cytochalasin D studies by disrupting the cytoplasmic actin network (Fig. 2D) rather by sequestering monomers and preventing their addition to actin filaments [11–13]. In this context, more cytosolic monomers would still be transferred to the nucleus but cannot participate in actin polymerisation. In stark contrast to Cytochalasin D treatment, nuclei in Latrunculin B-treated oocytes were not more likely to contain actin filaments (Fig. 2E) and only showed a two-fold increase in F-actin volume (Fig. 2F and S2A). Thus, the concentration of polymerisation-ready monomers transferred from the cytoplasm determines the degree of F-actin assembly in the oocyte nucleus. We directly tested whether high monomeric G-actin concentration is sufficient to induce nuclear F-actin polymerisation in oocytes. We exceeded endogenous nuclear monomeric G-actin concentration in mouse oocytes by targeting FLAG-beta-actin to the nucleus via the SV40 <u>n</u>uclear <u>l</u>ocalisation <u>s</u>ignal (NLS). This led to a ten-fold increase in nuclear F-actin volume in FLAG-beta-actin-NLS expressing oocytes, which were also more likely to contain F-actin than FLAG-NLS expressing Control oocytes (Fig. 2, G-I). Therefore, the concentration of monomeric G-actin in the nucleus indeed dictates the extent of nuclear F-actin polymerisation.

**Fig. 2.**
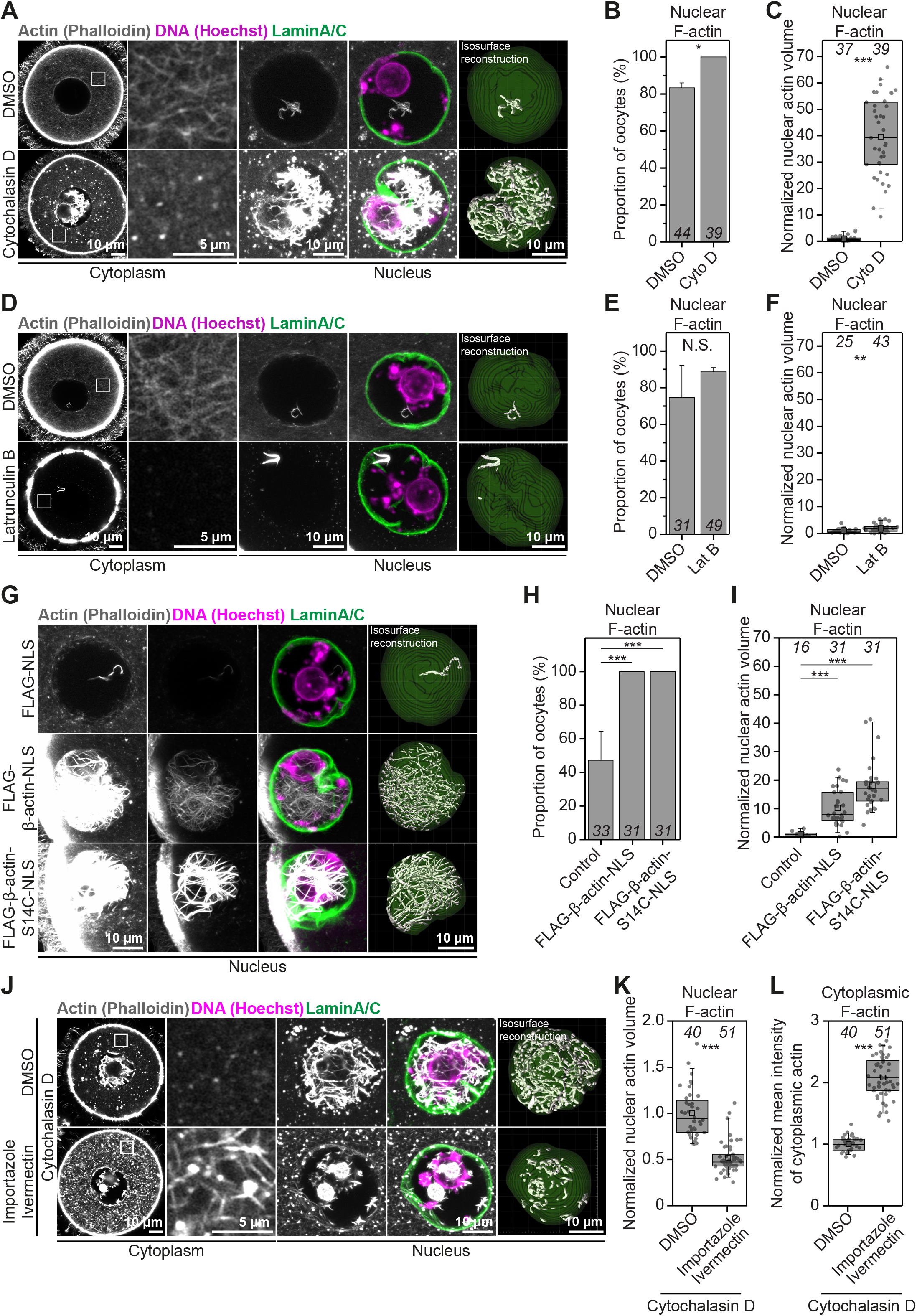
Excess G-actin causes uncontrolled cytoplasmic and nuclear F-actin assembly. (**A**) Single section Airyscan images of Phalloidin labelled cytoplasmic F-actin and maximum intensity projections (9 confocal sections) of nuclear actin filaments (grey), DNA (magenta) and nuclear membrane (green) in DMSO- or Cytochalasin D-treated mouse oocytes. Boxes in the oocyte cytoplasm mark regions that are magnified in insets. (**B**) Quantification of nuclear F-actin presence in DMSO- or Cytochalasin D-treated mouse oocytes. Data are from 3 independent experiments. (**C**) Quantification of nuclear F-actin volumes from isosurface reconstructions in A in DMSO- or Cytochalasin D-treated mouse oocytes. Data are from 3 independent experiments. (**D**) Single section Airyscan images of Phalloidin labelled cytoplasmic F-actin and maximum intensity projections (9 confocal sections) of nuclear actin filaments (grey), DNA (magenta) and nuclear membrane (green) in DMSO- or Latrunculin B-treated mouse oocytes. Boxes in the oocyte cytoplasm mark regions that are magnified in insets. (**E**) Quantification of nuclear F-actin presence in DMSO- or Latrunculin B-treated mouse oocytes. Data are from 3 independent experiments. (**F**) Quantification of nuclear F-actin volumes from isosurface reconstructions in D in DMSO- or Latrunculin B-treated mouse oocytes. Data are from 3 independent experiments. Y-axis scaling with higher resolution of data distribution in provided in Fig. S2A. (**G**) Maximum intensity projection (9 confocal sections) images of phalloidin labelled nuclear F-actin (grey), DNA (magenta) and nuclear membrane (green) in control and wild-type or S14C mutant actin expressing prophase-arrested oocytes. Excess nuclear F-actin in wild-type and S14C overexpressing oocytes is demonstrated by presenting overexposed (when nuclear F-actin is highly visible in controls) or moderately overexposed (when nuclear F-actin is poorly visible in controls) images. (**H**) Quantification of nuclear F-actin presence in control and wild-type or S14C actin mutant expressing mouse oocytes. Data are from 3 independent experiments. (**I**) Quantification of nuclear F-actin volumes from isosurface reconstructions in G in control and wild-type or S14C actin mutant expressing mouse oocytes. Data are from 3 independent experiments. (**J**) Single section Airyscan images of Phalloidin labelled cytoplasmic F-actin and maximum intensity projections (9 confocal sections) of nuclear actin filaments (grey), DNA (magenta) and nuclear membrane (green) in DMSO- or Importazole/Ivermectin-treated mouse oocytes that were then treated with Cytochalasin D. Boxes in the oocyte cytoplasm mark regions that are magnified in insets. (**K**) Quantification of nuclear F-actin volumes from isosurface reconstructions in J in DMSO- or Importazole/Ivermectin-treated mouse oocytes that were then treated with Cytochalasin D. Data are from 3 independent experiments. (**L**) Quantification of cytoplasmic F-actin network intensity in DMSO- or Importazole/Ivermectin-treated mouse oocytes that were then treated with Cytochalasin D. Data are from 3 independent experiments. Statistical significance was tested using Fisher’s exact test [(B), (E) and (H)] and Twotailed Student’s *t* test [(C), (F), (I), (K) and (L)].

To further examine the transfer of G-actin monomers to the nucleus, we blocked nuclear import in prophase-arrested oocytes using a combination of Importazole and Ivermectin [14, 15] (Fig. S3A), before disrupting the cytoplasmic actin network. Cytochalasin D treatment of oocytes did not cause excessive nuclear F-actin assembly when nuclear import was blocked (Fig. 2, C, J and K, S3B). In addition, nuclear import blockage caused distinct accumulation of actin on the surface of the oocyte nucleus (Fig. S3D), which supported the notion that actin monomers are nuclear import cargoes in oocytes. Surprisingly, Cytochalasin D treatment of nuclear import-defective oocytes resulted in a significantly denser cytoplasmic actin network that was composed of drug-resistant filaments (Fig. 2, J, L, S3C). This is consistent with a significant rise in cytosolic monomeric G-actin concentration, caused by blocked nuclear import, reducing Cytochalasin D activity [10]. Transfer of excess G-actin monomers to the nucleus is therefore necessary to maintain cytoplasmic F-actin network organisation in oocytes. This is supported by the observation that blocking nuclear import alone significantly increases cytoplasmic actin network density (Fig. S3, D and E). We propose that shuttling of cytosolic monomers into a large (~30 μm diameter) nucleus is a physiological G-actin buffering process that oocytes continuously use to modulate the cytoplasmic actin network.

To investigate meiotic consequences of dysfunctional G-actin buffering, we induced excess nuclear F-actin polymerisation by treating oocytes with Cytochalasin D (Fig. 2A) or by expressing a nuclear actin mutant (FLAG-beta-actin-S14C-NLS) that is more able to polymerise [16, 17] (Fig. 2, G-I). We then visualised chromatin (marked with SiR-DNA) and the nuclear envelope (marked with fluorescent nuclear membrane nanobodies) at high-temporal resolution. Initial analysis of these data indicated that excess nuclear actin filaments led to notably reduced chromatin mobility (Fig. 3, A and E, movies S3-S6), which in turn affects transcription in mouse oocytes [18]. We investigated this further by automated three-dimensional tracking of prominent chromatin spots throughout the nucleoplasm (Fig. 3, A and E). Indeed, when nuclei contained excess F-actin, chromatin spots showed significantly less movement over time (Fig. 3, B-D and F-H). Directly visualising chromatin (marked with histone H2B) and excess F-actin using overexpressed actin chromobody revealed that stable nuclear actin filaments can indeed physically entrap chromatin (Fig. S4A, movie S7).

Ultimately, excess nuclear F-actin bundles compromise meiosis and cause aneuploidy – these highly stable bundles persisted in FLAG-beta-actin-S14C-NLS expressing oocytes even after nuclear envelope disassembly (Fig. S4B) and severely interfered with chromosome alignment and segregation (Fig. 4, A-D and S4C, movies S8 and S9). Super-resolution immunofluorescence microscopy further demonstrated that after nuclear envelope disassembly stable nuclear F-actin structures become embedded in meiotic spindles and spindle poles where they obstruct chromosomal organisation (Fig. S4B). Therefore, homeostatic G-actin buffering in prophase oocytes must be fine-tuned to prevent excessive assembly of nuclear actin filaments that might interfere with successful completion of meiosis.

**Fig. 3.**
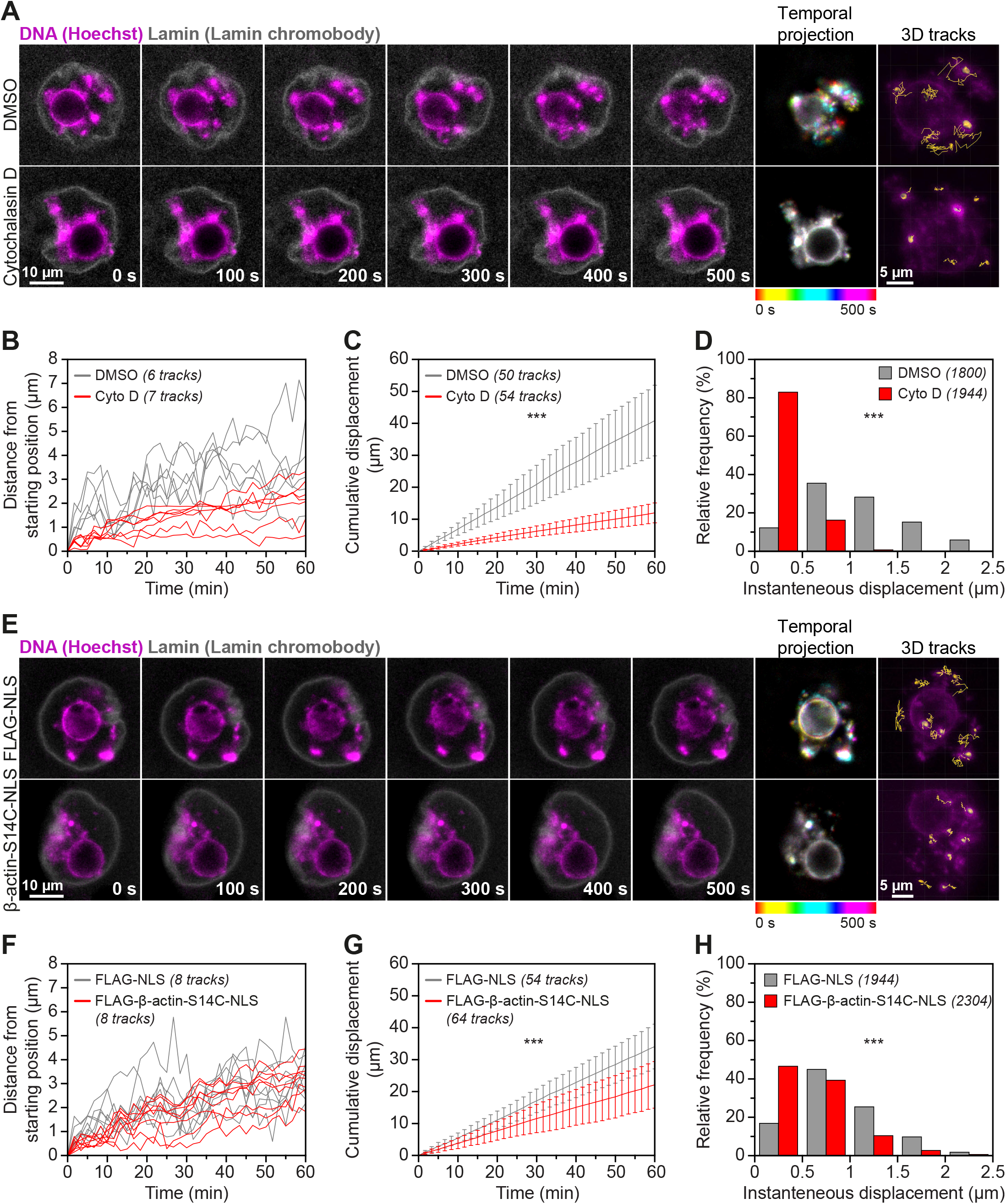
Excess nuclear actin filaments severely restrict oocyte chromatin mobility. (**A**) Stills from representative time lapse movies of chromatin movement in DMSO- or Cytochalasin D-treated mouse oocytes. Chromatin (SiR-DNA) is shown in magenta and nuclear membrane (Lamin chromobody) is shown in grey. Color-coded temporal projection images indicate the degree of chromatin mobility in the 500 seconds observation time. 3D tracks represent the spatial coverage of prominent chromatin spots over a 60 minutes observation period. (**B**) Distance from starting position of prominent chromatin spots in three dimensions over a 60-minute observation period in DMSO- or Cytochalasin D-treated mouse oocytes. Data are from 3 independent experiments. (**C**) Cumulative instantaneous displacement of prominent chromatin spots in three dimensions over a 60-minute observation period in DMSO- or Cytochalasin D-treated mouse oocytes. Data are from 3 independent experiments. Two-way analysis of variance was used to test for significance. (**D**) Relative frequencies of chromatin spot instantaneous displacement in DMSO- or Cytochalasin D-treated mouse oocytes. Data are from 3 independent experiments. Two-tailed Student’s *t* test was used to test for significance. (**E**) Stills from representative time lapse movies of chromatin movement in control or S14C actin mutant expressing mouse oocytes. Chromatin (SiR-DNA) is shown in magenta and nuclear membrane (Lamin chromobody) is shown in grey. Color-coded temporal projection images indicate the degree of chromatin mobility in the 500 seconds observation time. 3D tracks represent the spatial coverage of prominent chromatin spots over a 60 minutes observation period. (**F**) Distance from starting position of prominent chromatin spots in three dimensions over a 60-minute observation period in control or S14C actin mutant expressing mouse oocytes. Data are from 3 independent experiments. (**G**) Cumulative instantaneous displacement of prominent chromatin spots in three dimensions over a 60-minute observation period in control or S14C actin mutant expressing mouse oocytes. Data are from 3 independent experiments. Two-way analysis of variance was used to test for significance. (**H**) Relative frequencies of chromatin spot instantaneous displacement in control or S14C actin mutant expressing mouse oocytes. Data are from 3 independent experiments. Two-tailed Student’s *t* test was used to test for significance.

**Fig. 4.**
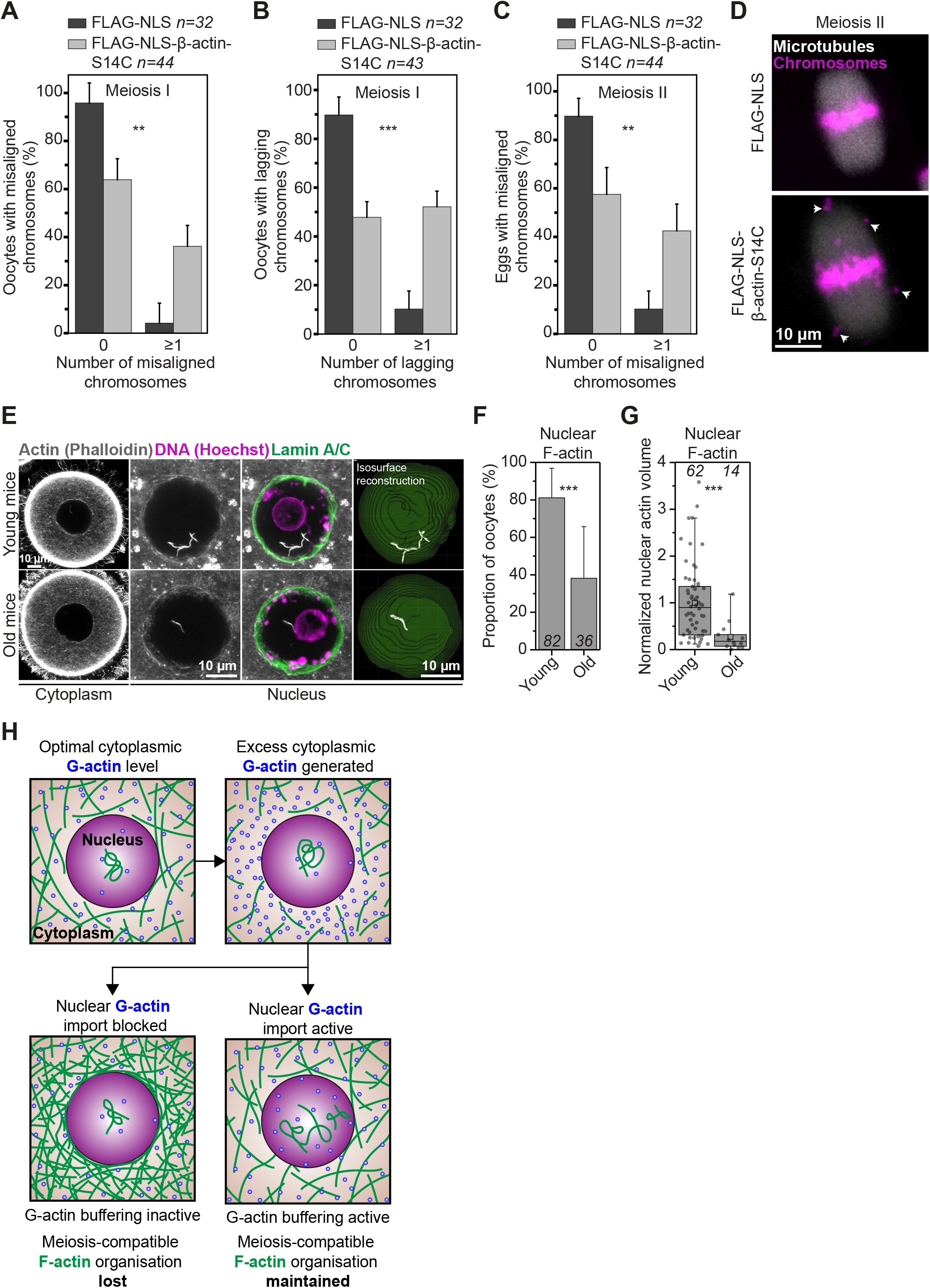
A defective homeostatic G-actin buffer contributes to mammalian oocyte aneuploidy. (**A**) Frequency of misaligned chromosomes in control (optimal nuclear F-actin in prophase) and S14C actin mutant expressing (excess nuclear F-actin in prophase) mouse oocytes. Data are from 4 independent experiments. (**B**) Frequency of lagging chromosomes in control (optimal nuclear F-actin in prophase) and S14C actin mutant expressing (excess nuclear F-actin in prophase) mouse oocytes. Data are from 4 independent experiments. (**C**) Frequency of misaligned chromosomes in control (optimal nuclear F-actin in prophase) and S14C actin mutant expressing (excess nuclear F-actin in prophase) mouse eggs. Data are from 4 independent experiments. (**D**) Representative images of fully aligned chromosomes in control (FLAG-NLS) and severely misaligned chromosomes (white arrows) in S14C actin mutant expressing mouse eggs. Microtubules (EGFP-MAP4-MTBD) are shown in grey and chromosomes (H2B-mRFP) are shown in magenta. (**E**) Single section Airyscan images of Phalloidin labelled cytoplasmic F-actin and maximum intensity projections (9 confocal sections) of nuclear actin filaments (grey), DNA (magenta) and nuclear membrane (green) in oocytes isolated from young and old mice. (**F**) Quantification of nuclear F-actin presence in oocytes isolated from young and old mice. Data are from 3 independent experiments. (**G**) Quantification of nuclear F-actin volumes from isosurface reconstructions in E in oocytes isolated from young and old mice. Data are from 3 independent experiments. (**H**) Model for regulation of cytoplasmic F-actin organisation using a homeostatic G-actin buffer. When G-actin buffering is active, excess cytosolic G-actin monomers are promptly transferred into the large oocyte nucleus. When it is defective, cytosolic G-actin concentration rises. This leads to assembly of a dense cytoplasmic actin network that interferes with the formation of healthy eggs. Statistical significance was tested using Fisher’s exact test [(A-C) and (F)] and Twotailed Student’s *t* test (G).

When prophase-arrested oocytes resume meiosis and initiate nuclear envelope disassembly en route to becoming eggs, density of the cytoplasmic F-actin network is progressively reduced – this is thought to support asymmetric cell division [4]. During this process, mouse oocytes were shown to control F-actin network density by adjusting the number and volume of vesicles in the cytoplasm [4]. In this context, homeostatic buffering of monomeric G-actin by the oocyte nucleus may aid vesicle-based actin network regulation.

Importantly, we found that the amount and complexity of nuclear F-actin structures declines significantly with increasing maternal age in non-manipulated mouse oocytes, with oocytes from 12 month old mothers having only 27% of the nuclear F-actin level observed in younger (8-12 week old) mothers (Fig. 4, E-G and Fig. S5, A and B). While we cannot fully exclude maternal age-dependent changes in cytosolic G-actin may contribute to nuclear F-actin decline, we did not detect changes in the cytoplasmic F-actin network of oocytes from older mothers (Fig. S5C). This raises the intriguing possibility that G-actin buffering defects may contribute to lower quality of eggs in reproductively older women.

Homeostatic G-actin buffering (Fig. 4H) could be a widely conserved function of large mammalian oocyte nuclei. For instance, prophase nuclei in non-manipulated sheep oocytes can also contain prominent nuclear actin filaments (Fig. S1C). Interestingly, nuclear F-actin is known to assemble in a variety of cellular contexts in non-gamete cells and embryos [8, 19–24] with postulated functions ranging from DNA repair to chromatin organisation. It will be important to explore whether the oocyte G-actin buffering process we describe here is a universal feature of mammalian nuclei and non-mammalian models where F-actin structures are intimately associated with the nucleus [25, 26]. In addition, mechanotransduction of actin-based forces to the nucleus is known to modulate nuclear mechanics and function in health and disease [27–29]. However, a direct role of the nucleus itself in this process by regulating cytosolic G-actin concentration, and thus F-actin assembly and force generation, should now be considered. Finally, our data indicate that commonly used actin drugs unintendedly stabilise nuclear F-actin and significantly affect nuclear mechanics. This will have important implications for past and future studies of sub-cellular actin-based structures in single- and multi-nucleated cells.

## Acknowledgment

We thank Paul Martin, David Stephens and Mark Dodding for discussion and critical comments on this manuscript, and the Animal Services Unit of the University of Bristol for technical assistance. We are grateful to Grazvydas Lukinavicius (MPI-BPC) for sharing improved and healthier derivatives of SiR-DNA dyes for labelling oocyte chromatin. This work was supported by a Sir Henry Dale Fellowship jointly funded by the Wellcome Trust and the Royal Society [grant number 213470/Z/18/Z]. The authors declare no competing financial interests.

K.S. performed experiments, analysed data, prepared figures, and revised the manuscript. F.G. performed experiments and revised the manuscript. B.M. performed experiments, analysed data, prepared figures, wrote the manuscript, and supervised the study.

## Supplementary Materials

Materials and Methods

Fig S1-S5

References (30-32)

Movies S1-S9

## Materials and Methods

### Preparation and microinjection of mammalian oocytes

All mice were maintained in a specific pathogen-free environment according to UK Home Office regulations and the guidelines of the University of Bristol Animal Services Unit. Oocytes were isolated from ovaries of 8-12 week old (young CD-1 mice), 10-12 months old (old CD-1 mice) or 8-12 weeks old 129 S6/SvEvTac mice, cultured, and microinjected with 6-8 picolitres of *in vitro* transcribed mRNA as described in detail recently [30].

Ovine ovaries were obtained from a University of Bristol Veterinary School slaughterhouse and transported to the laboratory at 37°C in M2 medium. Oocytes covered with several layers of cumulus cells were collected from ovaries by aspiration with an 18-gauge needle and cultured in M2 medium supplemented with 750 μM N6,2’-O-Dibutyryladenosine 3’,5’-cyclic monophosphate sodium salt (dbcAMP) before fixation and processing.

### Generation of expression constructs and mRNA synthesis

To mark microtubules, EGFP variant of the microtubule binding domain of mouse MAP4 (MAP4-MTBD, 659-1125 aa) [31] was generated as described in [7]. To label chromosomes, the coding sequence of Histone H2B was obtained from mouse cDNA and transferred by Gibson assembly into the HindIII site of pmRFP-N3 using primers 5’GGACTCAGATCTCGAGCTCAATGCCTGAGCCTGCGAAG3’ and 5’CCGTCGACTGCAGAATTCGACTTGGAGCTGGTGTACTTGG3’. The fragment corresponding to H2B-mRFP was then transferred into the XhoI-NotI site of pGEMHE [32] to generate the final construct pGEM-H2B-mRFP. To label nuclear actin, fluorescent nuclear actin nanobody (nuclear actin chromobody) plasmid was purchased from ChromoTek and transferred into the HindIII-EcoRI site of pGEMHE. To label the nuclear envelope, fluorescent lamin nanobody (lamin chromobody) plasmid was purchased from ChromoTek and transferred into the NcoI-XbaI site of pGEMHE. Wild type and S14C actin mutant expression constructs were generated using a synthetic construct encoding the SV40 nuclear localization signal (NLS) (5’CCGCCTAAGAAAAAGCGGAAGGTG3’) fused to mouse beta-actin (NM_007393.5). pGEM-FLAG-NLS beta-actin was generated by PCR linearization of pGEMHE with primers 5’ AATTCTGCAGTCGACGGC3’ and 5’ CGAAGCTTGAGCTCGAGATC3’ and joining by Gibson assembly with NLS-Beta-Actin that was flanked by primers 5’GATCTCGAGCTCAAGCTTCGATG***GACTACAAGGACGACGACGACAAG***GGGC CGCCTAAG3’ and 5’GGGCCGTCGACTGCAGAATTTTAGAAGCACTTGCGGTG3’. The coding sequence of the FLAG peptide DYKDDDDK in shown in bold italicized text. pGEM-FLAG-NLS-beta-actin-S14C mutant was generated by site-directed mutagenesis using primers 5’GTCGTCGACAACGGCTGCGGCATGTGCAAAGCC3’ and 5’GGCTTTGCACATGCCGCAGCCGTTGTCGACGAC3’.

Capped mRNA was synthesized using T7 polymerase (mMessage mMachine kit, following the manufacturer’s instructions, Ambion). mRNA concentrations were determined measurement on Nanodrop spectrophotometer (Thermo Scientific).

### Confocal and super-resolution live imaging

Images were acquired with Zeiss LSM800 microscope at 37°C. Oocytes were imaged in M2 medium under mineral oil using a 40x C-Apochromat 1.2 NA water-immersion objective as described in more detail in [30]. Super-resolution time-lapse images were acquired using the Airyscan module on Zeiss LSM800 microscope and processed post-acquisition using ZEN2.

### Immunofluorescence microscopy

Mouse and ovine oocytes were fixed with 100 mM HEPES, 50 mM EGTA, 10 mM MgSO_4_, 2% formaldehyde (v/v), and 0.5% Triton X-100 (v/v) at 37°C for 25-30 minutes (mouse) or for 60 minutes after 10 seconds pre-permeabilization in 0.4% Triton X-100 (v/v) in water (ovine). Oocytes were extracted in PBS supplemented with 0.3% Triton X-100 (v/v) at 4°C overnight. Antibody, F-actin and chromosome staining were performed for 2-2.5 hours in PBS, 3% BSA (w/v), and 0.1% Triton X-100 (v/v) at room temperature. Nuclear envelope was stained using primary rabbit anti-Lamin A/C antibody (ab133256, Abcam; 1:500) and Alexa-Fluor-647-labelled secondary anti-rat (Molecular Probes 1:400) antibodies. F-actin was stained with Rhodamine or Alexa-488 phalloidin (Molecular Probes; 1:20). DNA was stained with 5 μg/ml Hoechst 33342 (Molecular Probes).

Confocal and Airyscan super-resolution images were acquired with Zeiss LSM800 confocal microscope equipped with a 40x C-Apochromat 1.2 NA water-immersion objective. Images in control and perturbation conditions were acquired with identical imaging conditions.

### Drug addition experiments

To disrupt actin, oocytes were treated for 1 hour with Cytochalasin D (C8273-1MG, Merck) at a final concentration of 5 μg/ml or Latrunculin B (428020-1MG, Merck) at a final concentration of 5 μM in M2 medium supplemented with dbcAMP. To block nuclear import, oocytes were treated with a combination of 100 μM Importazole (SML0341-5MG, Merck) and 30 μM Ivermectin (I8000010, Merck). These final concentrations were maintained in experiments where oocytes were simultaneously treated with Cytochalasin D, Importazole and Ivermectin. All drugs were dissolved in DMSO (D2650-5X5ML, Merck). Where DMSO was used as control, it was diluted identically in M2 medium supplemented with dbcAMP to corresponding experimental conditions.

### Chromosome alignment and segregation analysis

For chromosome alignment and segregation analyses, images were acquired at a temporal resolution of 5 minutes and with a Z-stack thickness of ~40 μm at 1.5 μm confocal sections. Chromosomes that were distinctly separate from the metaphase plate chromosome mass at the time of anaphase onset (shown in Fig. 4D) were scored as misaligned chromosomes. Chromosomes that fell behind the main mass of segregating chromosomes for at least 10 minutes after anaphase onset (shown in Fig. S4C) were scored as lagging chromosomes. For both quantifications, maximum intensity projections of only those metaphase spindles that were oriented parallel to the imaging plane at and during anaphase were analysed.

### Isosurface reconstruction and 3D volume quantification of nuclear actin filaments

For 3D volume quantification of nuclear actin filaments, confocal images of nuclei in fixed oocytes were typically acquired at a spatial resolution of 1 μm confocal sections covering a range of 45 μm. Isosurfaces corresponding to nuclear membranes were reconstructed in three-dimension using the Cell module of Imaris software (Bitplane) and immunofluorescence signal of nuclear envelope antibodies. The nuclear isosurface was used to mask F-actin signal and to remove cytoplasmic F-actin structures surrounding the nuclei. In the masked region, three-dimensional isosurfaces of fluorescent phalloidin-labelled nuclear actin filaments were reconstructed using similar settings between different experimental groups within each repetition. Individual values for nuclear F-actin volume were normalized to the mean value of the control group for graphical presentation.

### Fluorescence intensity quantification of cytoplasmic F-actin

To quantify the density of the cytoplasmic actin network, single section super-resolution Airyscan images of fluorescent phalloidin-labelled cytoplasmic actin filaments were acquired. Mean fluorescence intensity of actin filaments was measured in the cytoplasm from six to twelve regions per oocyte and averaged to generate cytoplasmic F-actin intensity value for each oocyte. Background subtraction was performed in ImageJ by subtracting the mean fluorescence intensity of a region outside each oocyte from its corresponding cytoplasmic F-actin intensity value. Individual fluorescence intensity values were normalized to the mean value of the control group for graphical presentation.

### Four-dimensional tracking of prophase oocyte chromatin movement

For analyses of chromatin mobility, nuclear membranes in prophase-arrested oocytes were labeled by microinjecting and expressing fluorescent lamin nanobodies (lamin chromobody). In genetic nuclear actin stabilization experiments, FLAG-NLS or FLAG-NLS-beta-actin-S14C mRNA were co-expressed with nuclear lamin nanobodies. To label DNA, oocytes were incubated with 250 nM 5-SiR-Hoechst (SiR-DNA) in DMSO (Grazvydas Lukinavicius, MPI-BPC) for two hours before imaging experiment, during mRNA expression. In chemical nuclear actin stabilization experiments, oocytes were incubated in DMSO or Cytochalasin D one hour before imaging experiment, during mRNA expression. Confocal images of the nuclear envelope and chromosomes were acquired at a temporal resolution of 100 seconds with a Z-stack thickness of 36 μm at 1.5 μm confocal sections.

To exclude the contribution of nuclear movements to chromatin mobility, isosurfaces of the nuclear envelope were reconstructed using the Cell module of Imaris software (Bitplane) and fluorescent nuclear chromobody signal. Three-dimensional nuclear movement tracks obtained from these reconstructions were then used to automatically correct translational and rotational drift in Imaris. Isosurfaces of prominent chromatin spots were next reconstructed in three-dimensions using SiR-DNA fluorescence signal and tracked to automatically generate 3D tracks, which were manually corrected to remove inaccurate trajectories. Five to eight tracks of separate chromatin masses per oocyte were obtained and analyzed through this pipeline.

### Statistical data analyses

Histograms, statistical box plots and other graphs were generated using OriginPro software (OriginLab). Statistical box plots represent median (line), mean (small square), 5^th^, 95^th^ (whiskers) and 25^th^ and 75^th^ percentile (box enclosing 50% of the data) and are overlaid with individual data points. Average (mean), standard deviation and statistical significance based on two-tailed Student’s t test or Fisher’s exact test were calculated in OriginPro software (OriginLab). All error bars represent standard deviations. Two-way analysis of variance was performed in Prism software (GraphPad). Significance values are designated as * for *p* < 0.05, ** for *p* < 0.005 and *** for *p* < 0.0005. Non-significant values are indicated as N.S.

## Supplementary Movies

**Movie S1** Time lapse movie of nuclear F-actin (labelled with nuclear actin chromobody) in prophase-arrested mouse oocyte (oocyte 1)

**Movie S2** Time lapse movie of nuclear F-actin (labelled with nuclear actin chromobody) in prophase-arrested mouse oocyte (oocyte 2)

**Movie S3** Time lapse movie of chromatin movement in a DMSO-treated mouse oocyte. Chromatin (magenta) is labelled with SiR-DNA and nuclear membrane (grey) is labelled with lamin chromobody.

**Movie S4** Time lapse movie of chromatin movement in a Cytochalasin D-treated mouse oocyte. Chromatin (magenta) is labelled with SiR-DNA and nuclear membrane (grey) is labelled with lamin chromobody.

**Movie S5** Time lapse movie of chromatin movement in a FLAG-NLS expressing control mouse oocyte. Chromatin (magenta) is labelled with SiR-DNA and nuclear membrane (grey) is labelled with lamin chromobody.

**Movie S6** Time lapse movie of chromatin movement in a FLAG-NLS-beta-actin-S14C expressing mouse oocyte. Chromatin (magenta) is labelled with SiR-DNA and nuclear membrane (grey) is labelled with lamin chromobody.

**Movie S7** Time lapse movie of chromatin movement inside a mouse oocyte nucleus containing excess nuclear F-actin (induced by nuclear actin chromobody overexpression). Chromatin (magenta) is labelled with SiR-DNA and nuclear F-actin (green) is labelled with nuclear actin chromobody.

**Movie S8** Time lapse movie of chromosome alignment and segregation during meiosis I in a FLAG-NLS expressing control mouse oocyte. Microtubules (grey) are labelled with EGFP-MAP4-MTBD and chromosomes (magenta) are labelled with H2B-mRFP.

**Movie S9** Time lapse movie of chromosome alignment and segregation during meiosis I in a FLAG-NLS-beta-actin-S14C expressing mouse oocyte. Microtubules (grey) are labelled with EGFP-MAP4-MTBD and chromosomes (magenta) are labelled with H2B-mRFP.

---

**Fig. S1.**
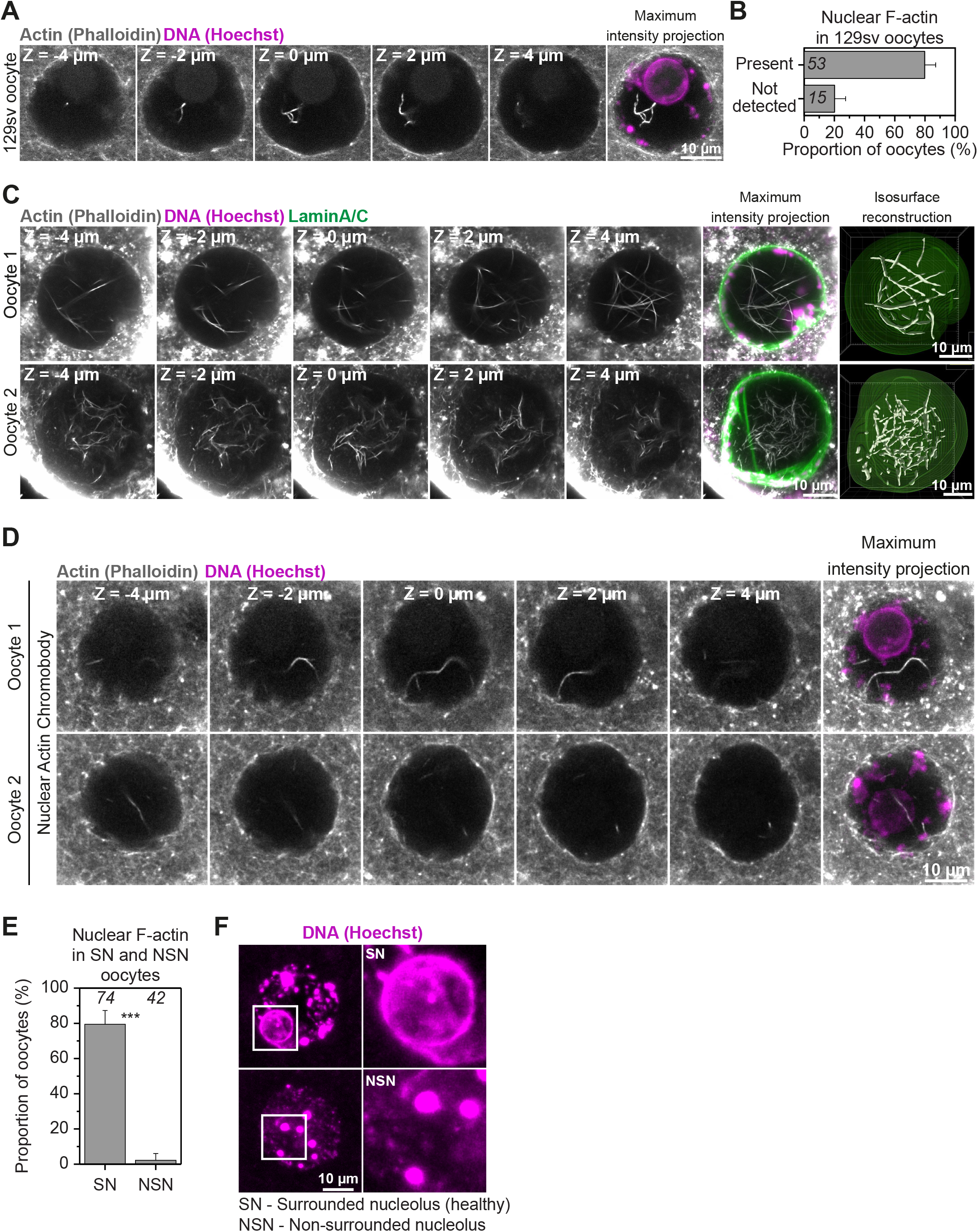
Nuclear F-actin is a common feature in healthy mammalian oocytes. (**A**) Phalloidin labelled nuclear actin filaments (grey) and chromosomes (Hoechst, magenta) in prophase-arrested oocyte isolated from 129sv mouse (inbred) strain. Single confocal sections spaced 2 μm apart and corresponding maximum intensity projections are shown. (**B**) Quantification of nuclear F-actin presence in prophase-arrested 129sv strain mouse oocytes. Data are from 3 independent experiments. (**C**) Maximum intensity projection (9 confocal sections) images of phalloidin labelled nuclear F-actin (grey), DNA (magenta) and nuclear membrane (green) in two sheep oocytes. Single confocal sections spaced 2 μm apart are shown. Isosurface reconstruction of actin (white) demonstrates prominent nuclear actin filaments. Sheep oocytes from two independent experiments are shown. (**D**) Phalloidin labelled nuclear actin filaments (grey) and chromosomes (Hoechst, magenta) in prophase-arrested mouse oocytes fixed after expression and live imaging of nuclear actin chromobody. Single confocal sections spaced 2 μm apart and corresponding maximum intensity projections are shown. (**E**) Quantification of nuclear F-actin presence in prophase-arrested mouse oocytes with surrounded nucleolar (SN) and non-surrounded nucleolar (NSN) chromatin configuration. Data are from 3 independent experiments. Fisher’s exact test was used to test for significance. (**F**) Representative images of surrounded nucleolar (SN) and non-surrounded nucleolar (NSN) chromatin (magenta) configuration in prophase-arrested oocytes. Boxes mark regions that are magnified in insets.

**Fig. S2.**
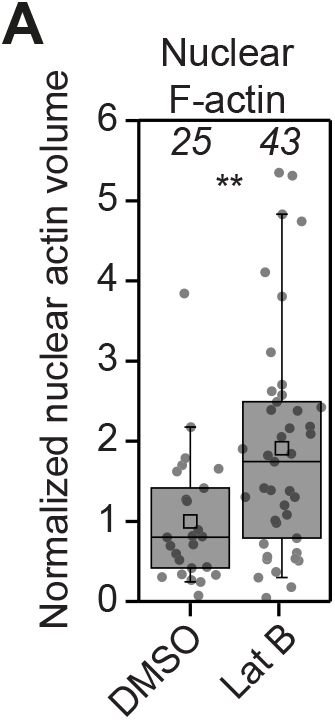
Excess cytosolic G-actin causes uncontrolled nuclear F-actin assembly. (A) Quantification of nuclear F-actin volumes from isosurface reconstructions in 2D in DMSO- or Latrunculin B-treated mouse oocytes. Data are from 3 independent experiments. Y-axis scaling is adjusted to show higher resolution distribution of data shown in Fig. 2F. Two-tailed Student’s *t* test was used to test for significance.

**Fig. S3.**
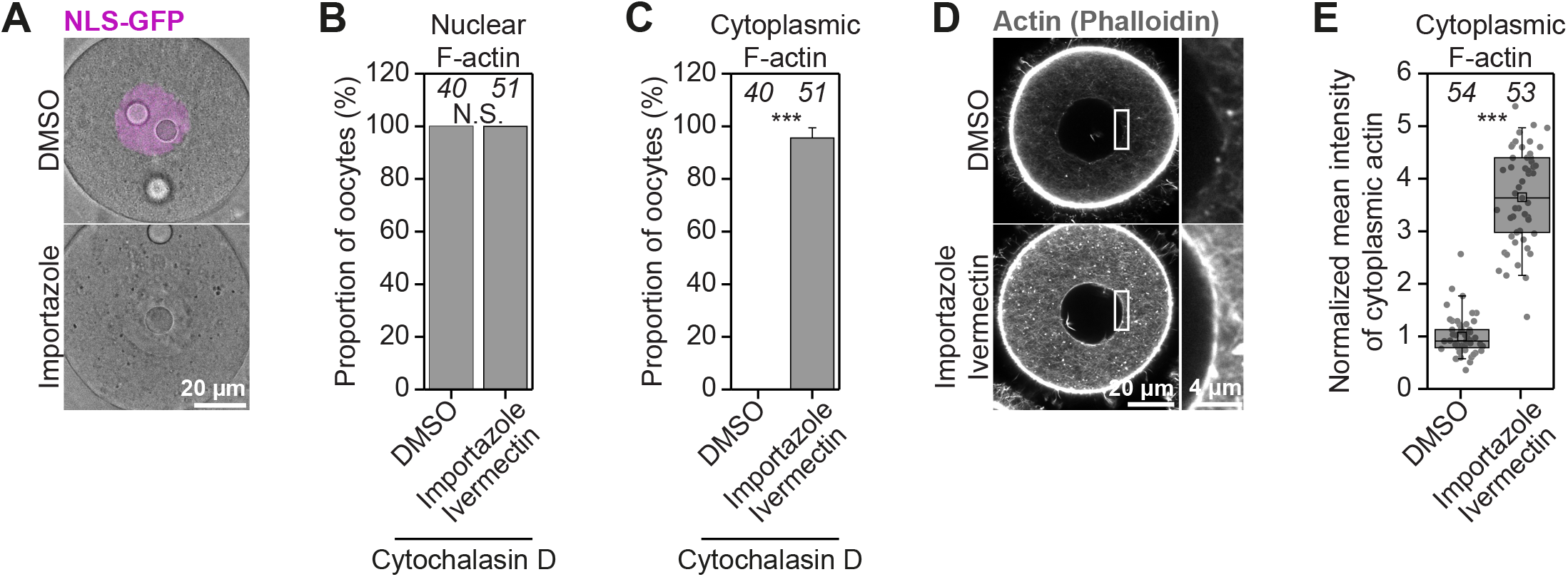
Actin monomers are nuclear import cargoes in mouse oocytes. (**A**) Representative images of GFP-NLS in DMSO- or Importazole-treated mouse oocytes. (**B**) Quantification of nuclear F-actin presence in DMSO- or Importazole/Ivermectin-treated mouse oocytes that were then treated with Cytochalasin D. Data are from 3 independent experiments. (**C**) Quantification of cytoplasmic F-actin network presence in DMSO- or Importazole/Ivermectin-treated mouse oocytes that were then treated with Cytochalasin D. Data are from 3 independent experiments. (**D**) Single section Airyscan images of Phalloidin labelled cytoplasmic F-actin in DMSO- or Importazole/Ivermectin-treated mouse oocytes. Boxes mark regions that are magnified in insets. (**E**) Quantification of cytoplasmic F-actin network fluorescence intensity in DMSO- or Importazole/Ivermectin-treated mouse oocytes. Data are from 3 independent experiments. Statistical significance was tested using Fisher’s exact test [(B) and (C)] and Twotailed Student’s *t* test (E).

**Fig. S4.**
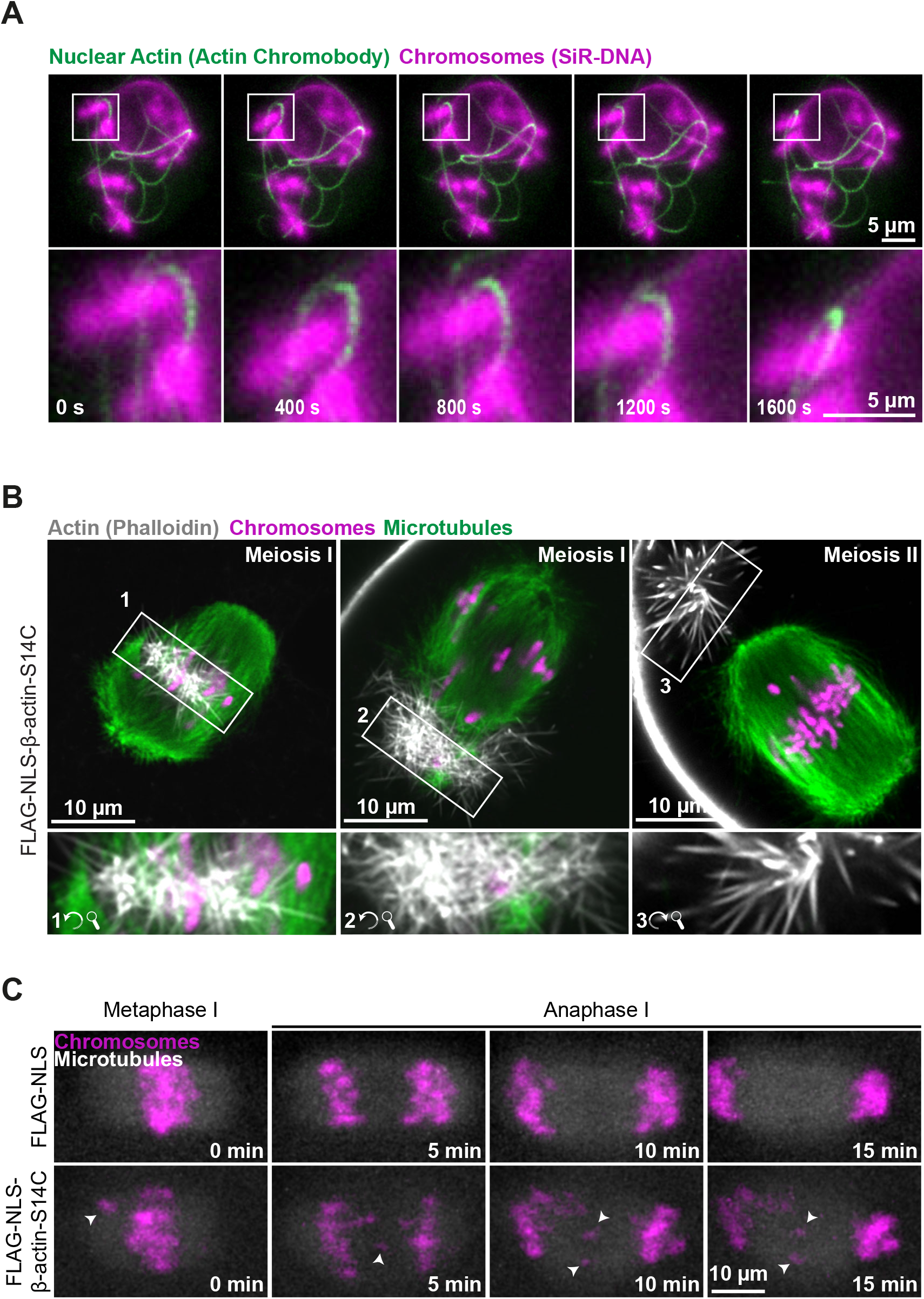
Excess nuclear F-actin compromises mammalian oocyte meiosis. (**A**) Stills from time lapse movie of chromatin (SiR-DNA, magenta) and nuclear actin (nuclear actin chromobody, green). Boxes mark regions magnified in insets and show entrapment of chromatin by actin filaments. (**B**) Single section Airyscan images of actin (grey), microtubules (green) and chromosomes (magenta) in S14C actin mutant expressing (excess nuclear F-actin in prophase) meiosis I oocytes and a meiosis II egg. Numbered boxes mark regions containing stable nuclear F-actin remnants that are rotated and magnified in insets. (**C**) Stills from representative time lapse movies of anaphase I in control or S14C actin mutant expressing oocytes. Microtubules (EGFP-MAP4-MTBD) are shown in grey and chromosomes (H2B-mRFP) are shown in magenta.

**Fig. S5.**
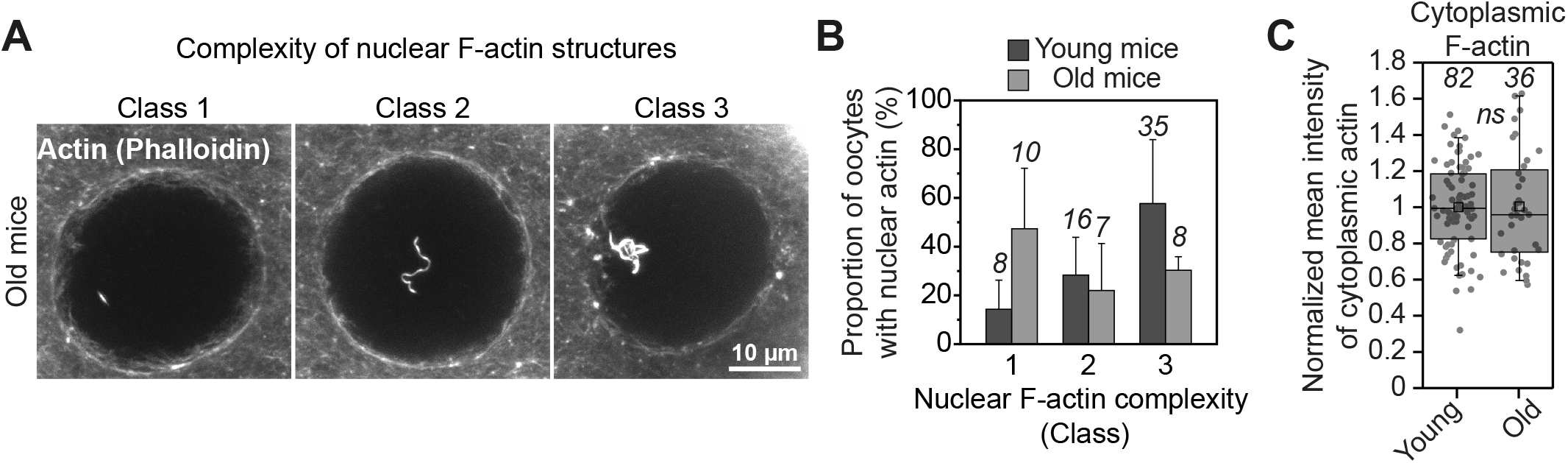
Nuclear F-actin declines with maternal age. (**A**) Three representative classes of nuclear F-actin (grey) complexity in phalloidin labelled mouse oocytes. (**B**) Quantification of the different classes of nuclear F-actin complexity (shown in A) in oocytes isolated from young and old mice. Data are from 3 independent experiments. (**C**) Quantification of cytoplasmic F-actin network intensity in oocytes from young and old mice. Data are from 3 independent experiments. Two-tailed Student’s *t* test.

## References

1. Mogessie, B., Scheffler, K., and Schuh, M. (2018). Assembly and Positioning of the Oocyte Meiotic Spindle. Annu Rev Cell Dev Biol.

2. Herbert, M., Kalleas, D., Cooney, D., Lamb, M., and Lister, L. (2015). Meiosis and maternal aging: insights from aneuploid oocytes and trisomy births. Cold Spring Harb Perspect Biol 7, a017970.

3. Schuh, M. (2011). An actin-dependent mechanism for long-range vesicle transport. Nat Cell Biol 13, 1431–1436.

4. Holubcova, Z., Howard, G., and Schuh, M. (2013). Vesicles modulate an actin network for asymmetric spindle positioning. Nat Cell Biol 15, 937–947.

5. Schuh, M., and Ellenberg, J. (2008). A new model for asymmetric spindle positioning in mouse oocytes. Curr Biol 18, 1986–1992.

6. Azoury, J., Lee, K.W., Georget, V., Rassinier, P., Leader, B., and Verlhac, M.H. (2008). Spindle positioning in mouse oocytes relies on a dynamic meshwork of actin filaments. Curr Biol 18, 1514–1519.

7. Mogessie, B., and Schuh, M. (2017). Actin protects mammalian eggs against chromosome segregation errors. Science 357.

8. Baarlink, C., Plessner, M., Sherrard, A., Morita, K., Misu, S., Virant, D., Kleinschnitz, E.M., Harniman, R., Alibhai, D., Baumeister, S., et al. (2017). A transient pool of nuclear F-actin at mitotic exit controls chromatin organization. Nat Cell Biol 19, 1389–1399.

9. Zuccotti, M., Giorgi Rossi, P., Martinez, A., Garagna, S., Forabosco, A., and Redi, C.A. (1998). Meiotic and developmental competence of mouse antral oocytes. Biol Reprod 58, 700–704.

10. Carlier, M.F., Criquet, P., Pantaloni, D., and Korn, E.D. (1986). Interaction of cytochalasin D with actin filaments in the presence of ADP and ATP. J Biol Chem 261, 2041–2050.

11. Spector, I., Shochet, N.R., Kashman, Y., and Groweiss, A. (1983). Latrunculins: novel marine toxins that disrupt microfilament organization in cultured cells. Science 219, 493–495.

12. Coue, M., Brenner, S.L., Spector, I., and Korn, E.D. (1987). Inhibition of actin polymerization by latrunculin A. FEBS Lett 213, 316–318.

13. Morton, W.M., Ayscough, K.R., and McLaughlin, P.J. (2000). Latrunculin alters the actin-monomer subunit interface to prevent polymerization. Nat Cell Biol 2, 376–378.

14. Soderholm, J.F., Bird, S.L., Kalab, P., Sampathkumar, Y., Hasegawa, K., Uehara-Bingen, M., Weis, K., and Heald, R. (2011). Importazole, a small molecule inhibitor of the transport receptor importin-beta. ACS Chem Biol 6, 700–708.

15. Wagstaff, K.M., Sivakumaran, H., Heaton, S.M., Harrich, D., and Jans, D.A. (2012). Ivermectin is a specific inhibitor of importin alpha/beta-mediated nuclear import able to inhibit replication of HIV-1 and dengue virus. Biochem J 443, 851–856.

16. Posern, G., Miralles, F., Guettler, S., and Treisman, R. (2004). Mutant actins that stabilise F-actin use distinct mechanisms to activate the SRF coactivator MAL. EMBO J 23, 3973–3983.

17. Wei, M., Fan, X., Ding, M., Li, R., Shao, S., Hou, Y., Meng, S., Tang, F., Li, C., and Sun, Y. (2020). Nuclear actin regulates inducible transcription by enhancing RNA polymerase II clustering. Sci Adv 6, eaay6515.

18. Almonacid, M., Al Jord, A., El-Hayek, S., Othmani, A., Coulpier, F., Lemoine, S., Miyamoto, K., Grosse, R., Klein, C., Piolot, T., et al. (2019). Active Fluctuations of the Nuclear Envelope Shape the Transcriptional Dynamics in Oocytes. Dev Cell 51, 145–157 e110.

19. Baarlink, C., Wang, H., and Grosse, R. (2013). Nuclear actin network assembly by formins regulates the SRF coactivator MAL. Science 340, 864–867.

20. Caridi, C.P., Plessner, M., Grosse, R., and Chiolo, I. (2019). Nuclear actin filaments in DNA repair dynamics. Nat Cell Biol 21, 1068–1077.

21. Plessner, M., Melak, M., Chinchilla, P., Baarlink, C., and Grosse, R. (2015). Nuclear F-actin formation and reorganization upon cell spreading. J Biol Chem 290, 11209–11216.

22. Wang, Y., Sherrard, A., Zhao, B., Melak, M., Trautwein, J., Kleinschnitz, E.M., Tsopoulidis, N., Fackler, O.T., Schwan, C., and Grosse, R. (2019). GPCR-induced calcium transients trigger nuclear actin assembly for chromatin dynamics. Nat Commun 10, 5271.

23. Kelpsch, D.J., and Tootle, T.L. (2018). Nuclear Actin: From Discovery to Function. Anat Rec (Hoboken) 301, 1999–2013.

24. Wesolowska, N., and Lenart, P. (2015). Nuclear roles for actin. Chromosoma 124, 481–489.

25. Mori, M., Somogyi, K., Kondo, H., Monnier, N., Falk, H.J., Machado, P., Bathe, M., Nedelec, F., and Lenart, P. (2014). An Arp2/3 nucleated F-actin shell fragments nuclear membranes at nuclear envelope breakdown in starfish oocytes. Curr Biol 24, 1421–1428.

26. Wesolowska, N., Avilov, I., Machado, P., Geiss, C., Kondo, H., Mori, M., and Lenart, P. (2020). Actin assembly ruptures the nuclear envelope by prying the lamina away from nuclear pores and nuclear membranes in starfish oocytes. Elife 9.

27. Isermann, P., and Lammerding, J. (2013). Nuclear mechanics and mechanotransduction in health and disease. Curr Biol 23, R1113–1121.

28. Martino, F., Perestrelo, A.R., Vinarsky, V., Pagliari, S., and Forte, G. (2018). Cellular Mechanotransduction: From Tension to Function. Front Physiol 9, 824.

29. Alisafaei, F., Jokhun, D.S., Shivashankar, G.V., and Shenoy, V.B. (2019). Regulation of nuclear architecture, mechanics, and nucleocytoplasmic shuttling of epigenetic factors by cell geometric constraints. Proc Natl Acad Sci U S A 116, 13200–13209.

30. Mogessie, B. (2020). Visualization and Functional Analysis of Spindle Actin and Chromosome Segregation in Mammalian Oocytes. Methods Mol Biol 2101, 267–295.

31. Mogessie, B., Roth, D., Rahil, Z., and Straube, A. (2015). A novel isoform of MAP4 organises the paraxial microtubule array required for muscle cell differentiation. Elife 4, e05697.

32. Schuh, M., and Ellenberg, J. (2007). Self-organization of MTOCs replaces centrosome function during acentrosomal spindle assembly in live mouse oocytes. Cell 130, 484–498.

